## Supplementary material for "Natural xanthones as α-Mangostin induce vasorelaxation involving key gating residues in the S6 domain of BK channels": Source data file 1

**Source data file 1:** Summary of data used in figures and applied statistical tests with p-values.

**to**

Tables S1 - S16

*Table S1: Modulation of different $K^+$ Channels by α -Mangostin shown by current fold change ± SEM. To test for statistical significance of activation or inhibition, relative currents before and after α-Mangostin application (10 µM) were compared with multiple t-tests with the Holm-Sidak correction for multiple comparisons (α=0.05).*

| Fig. 1A/S1 | Channel | Fold Change ($I/I_0$) at +40 mV | Adjusted P value |
|---|---|---|---|
| | TREK-1 | 10.3 ± 1.64 | <0.001 |
| | TALK-1 | 0.9 ± 0.04 | 0.15 |
| | TRESK | 0.51 ± 0.12 | 0.009 |
| | TWIK-1$^m$ | 0.12 ± 0.03 | <0.001 |
| | TWIK-2$^m$ | 0.45 ± 0.06 | <0.001 |
| | TASK-3 | 0.2 ± 0.04 | <0.001 |
| | THIK-1 | 0.91 ± 0.05 | 0.15 |
| | BKα | 20.41 ± 2.39 | <0.001 |
| | BKα/β1 | 24.42 ± 3.85 | <0.001 |
| | $K_v$1.1 | 1.16 ± 0.13 | 0.25 |
| | $K_v$1.3 | 0.4 ± 0.04 | <0.001 |
| | $K_v$11.1 (hERG) | 0.84 ± 0.07 | 0.15 |
| | $K_{ir}$ 1.1 (ROMK) | 0.8 ± 0.09 | 0.15 |
| | $K_{ir}$ 2.1 | 0.68 ± 0.04 | <0.001 |

*Table S2: Current fold change ($I/I_0$) ± SEM for application of different Mangostin compounds/ formulations on TREK-1 and BK channels compared to α-Mangostin. One-way ANOVA with Dunnett's multiple comparisons post-hoc test for each group (α=0.05); for TREK-1 F=0.196, P=0.82; for BKα F=7.65, P=0.001; for BKα/β1 F=0.903, P=0.42.*

| Fig. 1D | Fold Change ($I/I_0$) at +40 mV | | | Adjusted P values | |
|---|---|---|---|---|---|
| | α-Mangostin | γ-Mangostin | Dietary suppl. | γ-Mangostin | Dietary suppl. |
| TREK-1 | 10.3 ± 1.64 | 11.35 ± 1 | 9.18 ± 2.74 | 0.87 | 0.94 |
| BKα | 20.41 ± 2.39 | 38.54 ± 4.8 | 19.98 ± 2.99 | 0.001 | >0.99 |
| BKα/β1 | 24.42 ± 3.85 | 29.32 ± 5.03 | 20.92 ± 3.49 | 0.66 | 0.83 |

*Table S3: Comparison of the $V_{½}$ ± SEM and slope ± SEM of the GV-relationship (Boltzmann fit) before and after activation by 10 µM α-Mangostin in BKα and BKα/β1 channels. $V_{½}$: paired t-test, two-tailed P value (α=0.05); slope: Wilcoxon matched-pairs signed rank test, two-tailed exact P value (α=0.05).*

| Fig. 2B | $V_{½}$ (mV) | | Δ $V_{½}$ (mV) | P value | slope (mV) | | P value |
|---|---|---|---|---|---|---|---|
| | basal | 10 µM α-Mangostin | | | basal | 10 µM α-Mangostin | |
| BKα | 110.49 ± 2.69 | 57.37 ± 3.60 | 53.08 ± 4.9 | <0.001 | 25.9 ± 1.33 | 21.8 ± 1.08 | 0.06 |
| BKα/β1 | 147.25 ± 5.66 | 64.83 ± 4.25 | 82.42 ± 4.96 | <0.001 | 28.8 ± 1.19 | 28.4 ± 3.24 | 0.84 |

*Table S4: Comparison of activation and deactivation time constants (τ) of BKα and BKα/β1 channels in the absence and after application of 10 µM α-Mangostin (mean ± SEM). Paired t-tests, two-tailed P values (α=0.05).*

| Fig. 2C | BKα | | BKα/β1 | |
|---|---|---|---|---|
| | Activation +100 mV | Deactivation +100 mV | Activation +100 mV | Deactivation +100 mV |
| basal | 7.96 ± 1.64 ms | 0.90 ± 0.04 ms | 63.76 ± 16.03 ms | 3.60 ± 0.16 ms |
| 10 µM α-Mangostin | 4.71 ± 0.71 ms | 6.85 ± 1.11 ms | 12.36 ± 1.20 ms | 95.6 ± 13.99 ms |
| P value | 0.02 | 0.002 | 0.02 | 0.001 |

*Table S5: Activation (n=7) and deactivation (n=6) time constants (τ) of BKα channels measured in 10 µM free $Ca_i^{2+}$ for example voltages in the physiological range in the absence and after application of 10 µM α-Mangostin (mean ± SEM). Paired t-tests, two-tailed P values (α=0.05).*

| **Fig. 2 S1** | **Activation τ** | **Deactivation τ** |
|---|---|---|
| | activation +20 mV | prepulse +20 mV |
| basal | 12.2 ± 0.6 ms | 1.02 ± 0.21 ms |
| 10 µM α-Mangostin | 4.98 ± 1.6 ms | 5.59 ± 1.03 ms |
| P value | 0.003 | 0.006 |

*Table S6: Shift of voltage activation ± SEM in BKα channels in different free $Ca^{2+}$ concentrations. Difference of shifts in $V_{½}$ ± SEM in higher $Ca^{2+}$ concentrations compared to 0.1 µM $Ca^{2+}$ were tested with the Kruskal-Wallis test (α=0.05).*

| **Fig. 2D** | **free $Ca^{2+}$ (µM)** | **$V_{½}$ (mV)** | **$V_{½}$ (mV) in 10 µM α-Mangostin** | **Δ $V_{½}$ (mV)** | **Exact P value** |
|---|---|---|---|---|---|
| | 0.1 | 110.49 ± 2.69 | 57.37 ± 3.60 | 53.08 ± 4.9 | - |
| | 1 | 91.17 ± 4.20 | 7.85 ± 6.6 | 82.74 ± 10.17 | 0.02 |
| | 10 | 13.9 ± 2.52 | -36.65 ± 8.13 | 50.55 ± 9.13 | >0.99 |

*Table S7: Single channel analysis of BKa channels measured in excised inside-out patches from HEK293 cells. Unpaired t-tests, two-tailed P values (α=0.05).*

| **Fig. 3B, C** | **Open probability** | **Amplitude (pA)** | **Open Dwell time (ms)** | **Closed Dwell time (ms)** |
|---|---|---|---|---|
| basal | 0.002 ± 0.0008 | 9.18 ± 0.29 | -0.84 ± 0.45 | 2.38 ± 0.23 |
| 10 µM α-Mangostin | 0.77 ± 0.08 | 9.76 ± 0.26 | 0.80 ± 0.24 | -1.39 ± 0.36 |
| P value | <0.001 | 0.18 | 0.002 | <0.001 |

*Table S8: Burst analysis of single BKa channels measured in excised inside-out patches from HEK293 cells in 5 µM free $Ca_i^{2+}$. Mean of means from Fig. 3E are presented (n=3 patches). Unpaired t-tests, two-tailed P values (α=0.05).*

| **Fig. 3E, F** | **Burst duration (ms)** | **Long closures (ms)** | **Openings/ burst** | **Intraburst closed time (ms)** | **Intraburst open time (ms)** |
|---|---|---|---|---|---|
| basal | 6383 ± 2095 | 7423 ± 1855 | 316.8 ± 62.5 | 20.79 ± 1.15 | 18.62 ± 1.23 |
| 10 µM α-Mangostin | 27429 ± 12017 | 3553 ± 1412 | 536.6 ± 159 | 29.44 ± 1.26 | 28.91 ± 1.18 |
| P value | n.d. | n.d. | n.d. | <0.001 | <0.001 |

*Table S9: TPA $IC_{50}$ ± SEM values in the absence and presence of α-Mangostin for the competition experiment in TREK-1 channels. An unpaired t-test was used to test for difference to TPA alone (two-tailed P value, α=0.05).*

| **Fig. 4 S1B** | **TPA $IC_{50}$ (µM)** | **P value** |
|---|---|---|
| TPA | 0.72 ± 0.64 | - |
| TPA + 5 µM α-Mangostin | 6.33 ± 0.86 | 0.002 |

*Table S10: Current fold change ± SEM ($I/I_0$) after application of 5 µM α-Mangostin to cysteine mutants in the M2 and M4 segment of TREK-1 channels. Brown-Forsythe and Welsh ANOVA with Welch's correction, α=0.05, F (DFn, DFd) = 10.68 (10, 35.84), W (DFn, DFd) = 12.19 (10, 30.22) was used to test for differences to the WT channel after α-Mangostin application.*

| **Fig. 4 S1C** | **Fold change ($I/I_0$)** | **P value** |
|---|---|---|
| TREK-1 WT | 5.36 ± 0.82 | - |
| G181C | 6.55 ± 2.03 | <0.001 |
| I182C | 1.68 ± 0.18 | 0.001 |
| P183C | 67.12 ± 14.66 | <0.001 |
| L184C | 3.79 ± 0.81 | 0.19 |
| V303C | 5.21 ± 0.84 | 0.90 |
| L304C | 2.49 ± 0.31 | 0.006 |
| S305C | 2.1 ± 0.35 | 0.003 |
| M306C | 2.31 ± 0.28 | 0.004 |
| I307C | 2.38 ± 0.5 | 0.007 |
| G308C | 1.02 ± 0.03 | <0.001 |

*Table S11: Comparison of estimated THexA $IC_{50}$ values from competition experiments with α-Mangostin or BC5 in BKα channels. Kruskal-Wallis test with Dunn's post-hoc test (α=0.05).*

| **Fig. 4A** | **THexA $IC_{50}$ of BKα channels (nM)** | **Adjusted P value** |
|---|---|---|
| THexA | 77.51 ± 5.53 | - |
| THexA + 10 µM α-Mangostin | 1642.29 ± 577.82 | 0.04 |
| THexA + 100 µM BC5 | 45.6 ± 8.05 | 0.31 |

*Table S12: Shifts in half-maximal activation voltage ($V_{½}$) in the BKa WT channel for 10 µM α-Mangostin applied at different pH. Brown-Forsythe and Welch ANOVA with Dunnet's T3 multiple comparison post-hoc test (α=0.05), F (DFn, DFd) = 25.7 (2.00, 10.7); W (DFn, DFd) = 28.9 (2.00, 8.58).*

| **Fig. 4C, S2** | **$V_{½}$ (mV)** | **$V_{½}$ (mV) in 10 µM α-Mangostin** | **Δ $V_{½}$ (mV)** | **Adjusted P value** |
|---|---|---|---|---|
| pH 8.5 | 91.49 ± 3.04 | 56.77 ± 4.06 | 75.97 ± 2.02 | 0.004 |
| pH 7.2 | 109.56 ± 3.02 | 55.51 ± 3.23 | 54.05 ± 4.04 | - |
| pH 6 | 127.61 ± 3.05 | 51.64 ± 4.23 | 34.71 ± 5.63 | 0.04 |

*Table S13: Voltages of half-maximal activation ($V_{½}$) before and after activation with 10 µM α-Mangostin and the resulting shift in $V_{½}$ for WT different mutant BKα channels. Brown-Forsythe and Welsh ANOVA with Dunnet's T3 post-hoc test (α=0.05), F (DFn, DFd) = 16.72 (6, 30.4), W (DFn, DFd) = 30.84 (6, 17.64).*

| **Fig. 4D, E** | **$V_{½}$ (mV)** | **$V_{½}$ (mV) in 10 µM α-Mangostin** | **Δ $V_{½}$ (mV)** | **Adjusted P value** |
|---|---|---|---|---|
| BKα WT | 110.45 ± 2.69 | 57.37 ± 3.60 | 53.08 ± 4.9 | - |
| I308A | 54.45 ± 3.86 | 34.48 ± 3.81 | 19.97 ± 3.12 | <0.001 |
| L312M | 126.29 ± 6.19 | 98.39 ± 4.46 | 27.89 ± 5.42 | 0.02 |
| A316P | 85.6 ± 2.71 | 81.04 ± 3.58 | 4.56 ± 1.23 | <0.001 |
| A316G | 36.7 ± 2.96 | 13.13 ± 3.11 | 23.57 ± 2.05 | 0.002 |
| S317R | 120.57 ± 0.96 | 86.24 ± 3.41 | 34.33 ± 3.37 | 0.04 |
| Y318S | 112.87 ± 3.25 | 71.41 ± 5.63 | 41.46 ± 5.17 | 0.52 |

*Table S14: Shifts in half-maximal activation voltage ($V_{½}$) for 1 µM GoSlo-SR-5-6 in the BKa WT channel and I308A and A316P mutants. One-way ANOVA with Dunnet's multiple comparison post-hoc test (α=0.05), F=6.51, F (DFn, DFd) = 2.09 (2, 17).*

| **Fig. 4F** | **Channel** | **$V_{½}$ (mV)** | **$V_{½}$ (mV) activated** | **Δ $V_{½}$ (mV)** | **Adjusted P value** |
|---|---|---|---|---|---|
| GoSlo-SR-5-6 (1 µM) | BKα WT | 104.17 ± 5.85 | 61.46 ± 6.35 | 42.71 ± 3.05 | - |
| | A316P | 78.25 ± 6.92 | 50.86 ± 5.53 | 25.0 ± 2.81 | 0.01 |
| | I308A | 53.83 ± 1.27 | 27.94 ± 4.68 | 25.89 ± 4.05 | 0.008 |

*Table S15: Vehicle control (0.025 % DMSO) for BKα (n=4) and $Ca_v1.2$ channels (n=3) as mean ± SEM. Paired t-tests, two-tailed P values (α=0.05).*

| Fig. 5 S1 | BKα | | $Ca_v1.2$ | |
|---|---|---|---|---|
| | $V_{½}$ (mV) | $V_{½}$ (mV) DMSO | Norm. current at 0 mV | Norm. current at 0 mV (DMSO) |
| | 66.3 ± 6.1 | 68.8 ± 5.6 | -0.94 ± 0.06 | -0.95 ± 0.03 |
| P-value | | 0.45 | | 0.92 |

*Table S16: Contraction force of aortic preparations after application of 10 µM α-Mangostin. Kruskal-Wallis test with Dunn's post-hoc test for multiple comparison (α=0.05).*

| Fig. 5C | norm. contraction force ($f/f_{NA}$) | | Adjusted P value |
|---|---|---|---|
| | mean ± SEM | median | |
| 10 µM α-Mangostin | 0.16 ± 0.08 | 0.113 | <0.001 |
| 100 nM IbTx | 1.04 ± 0.02 | 1.05 | - |
| 100 nM IbTx + 10 µM α-Mangostin | 0.78 ± 0.12 | 1.02 | 0.74 |
